# Amino acids activate parallel chemosensory pathways in *Drosophila*

**DOI:** 10.1101/2025.03.10.642418

**Authors:** Jacqueline Guillemin, Grace Davis, Kayla Audette, Tucker Avonda, Ella Freed, Ava Vitters, Braden Woods, Erinn Wagner, Lauren Schwartz, Ian Orsmond, Beckett Hampp, Megan Burdick, Peter Gause, Sascha Taylor, Brenna Asaro, Alice Sperber, Kaitlyn A. Zoller, Molly Stanley

**Author notes:** co-first authors (Jacqueline Guillemin, Grace Davis).

## Abstract

Amino acids (AAs) are essential dietary macronutrients that impact an organism’s fitness in a concentration-dependent manner, but the mechanisms mediating AA detection to drive consumption are less clear. In *Drosophila*, we identified the full repertoire of taste cells and receptors involved in feeding initiation towards a glutamate-rich AA mixture, tryptone, using *in vivo* calcium imaging and the proboscis extension response (PER). We found that AA attraction occurs through sweet cells, whereas feeding aversion is mediated through Ionotropic Receptor 94e (IR94e) cells and bitter cells, dependent on concentration. Further, our results corroborate previous findings that ionotropic receptors *IR76b*, *IR51b*, and *IR94e* detect AAs in their respective cell types. Additionally, we describe a new role for the appetitive IR56d receptor and bitter gustatory receptors in detecting AAs. This work establishes a cellular and molecular framework of AA feeding initiation and highlights redundancy in aversive pathways that regulate AA feeding.

## INTRODUCTION

Consuming energy-dense nutrients is vital for survival^1^. Protein is a dietary source of amino acids (AAs), which are essential for optimal health but harmful in excess^2–4^. In addition, protein needs vary significantly based on internal states, such as for reproduction^5,6^. Animals must detect AAs in a way that allows for flexible behaviors, but the gustatory mechanisms used to sense proteinogenic AAs to guide feeding are still under investigation. Glutamate, the AA behind the ‘umami’ flavor, is the most abundant AA in foodstuff^7^ and has been the focus of studies of mammalian taste where an AA taste receptor was identified^8,9^. However, individual AAs have unique tastes to humans^10–12^, and there is evidence that AAs can also activate “sweet” or “bitter” taste receptors^8,9,13,14^.

Investigations into the neural circuits encoding AA taste are attainable in the fruit fly, *Drosophila melanogaster*, where genetic and neurobiological tools have led to the development of single-cell mapping and a full-brain connectome^15–17^. Previous work indicates that certain individual AAs are detected by chemosensory ionotropic receptors (IRs) or gustatory receptors (GRs) in *Gr64f-*expressing “sweet” cells and/or *Gr66a-*expressing “bitter” cells^18,19^. Additionally, we recently identified a set of gustatory receptor neurons (GRNs) on the labellum of *Drosophila* that express *Ionotropic Receptor 94e* (*IR94e)* and detect the AA glutamate^20^. If and how all three cell types potentially work in parallel to detect AAs and promote or discourage feeding is currently unclear.

This report identifies the full repertoire of GRNs and receptors that are salient for feeding initiation when flies encounter proteinogenic AAs in a food source. Utilizing the proboscis extension response (PER) and *in vivo* calcium imaging, we found that flies initiate feeding to tryptone, a physiological AA mixture with relatively high levels of glutamate, through the activation of sweet, IR94e, and bitter GRNs via IRs and GRs. Notably, some of the receptors found here contrast with those identified for single AAs^18^, suggesting that a mix of AAs may interact uniquely with receptors on the labellum. These findings illustrate the necessity of a combinatorial coding system in which AAs in foodstuffs activate multiple neural circuits to provide complex sensory signals to promote fitness.

## RESULTS

### A mixture of amino acids drives feeding initiation via the labellum

To identify a solution containing proteinogenic AAs in ratios similar to natural foodstuff without other tastants, we assessed the AA composition of four food sources common to *Drosophila melanogaster*^21^: apple^22–25^, banana^23,25,26^, grape^22,23,25^, and tomato^23^. Composition data from the United States Department of Agriculture delineated aspartic or glutamic acid as the most concentrated AA in these fruits, while the other 18 proteinogenic AAs varied in concentration (Fig. 1A)^27^. Yeast is a natural source of protein for *Drosophila,* and previous studies have used whole yeast or yeast extract to study AA taste and feeding behaviors^28–32^. However, these solutions contain other molecules, such as ribonucleotides, metals, B vitamins, and glycerol, which could independently impact taste behaviors^33–36^. Regarding the AA composition of yeast extract, alanine is the most abundant, with lower relative concentrations of aspartic and glutamic acids (Fig. 1A)^33^. Tryptone, a casein digest, has been used as a yeast alternative to study protein satiety in *Drosophila*^37^. Glutamic acid is the most concentrated AA in tryptone, similar to the aforementioned fruits (Fig. 1A)^38^. Therefore, we used tryptone to study the salient detection mechanisms that drive feeding initiation when flies encounter a mix of AAs in physiologically relevant ratios.

**Figure 1:**
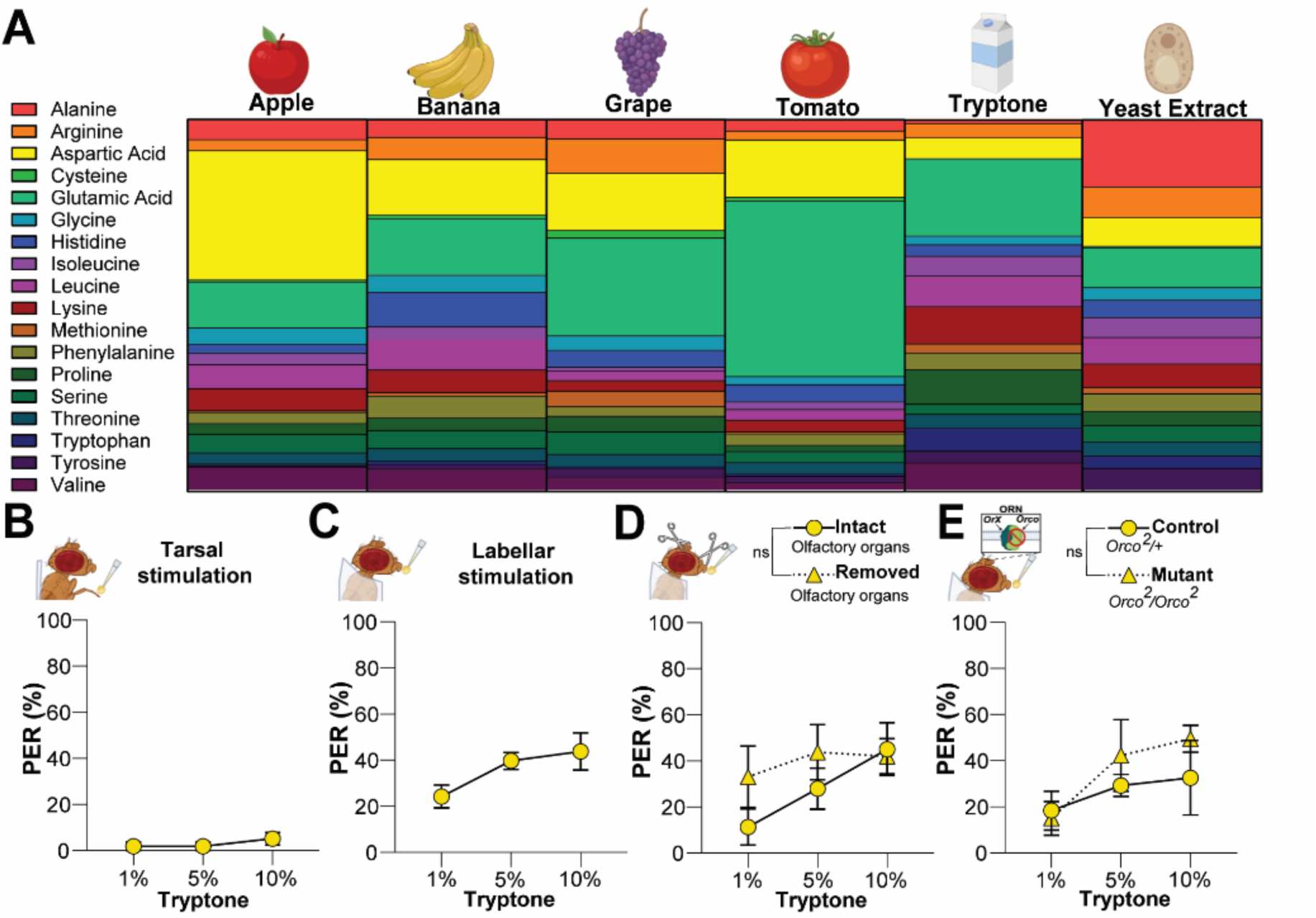
The labellum detects a physiological mix of amino acids to initiate feeding. **(A)** AA breakdown of prevalent *Drosophila* foodstuffs compared to tryptone and yeast extracts. **(B)** Tarsal and (**C**) Labellar PER to a tryptone concentration gradient, n=11-12 groups of 7-10 flies. (**D**) Labellar PER to tryptone following removal of the antennae and maxillary palps, n=5 groups of 7-10 flies. (**E**) Labellar PER to tryptone in *Orco* mutants, n=5 groups of 7-10 flies. ns=p>0.05 for main effect of genotype by two-way ANOVA. Each graph depicts mean ± SEM.

To establish if flies initiate feeding to tryptone, we performed a dose-response curve using the proboscis extension response (PER)^39^. All experiments were performed on mated females, as they have a strong drive to consume AAs^30,31^. PER occurs when attractive gustatory receptor neurons (GRNs) on the legs or the labellum are sufficiently activated relative to any co-activation of aversive GRNs^39,40^. Tarsal stimulation with tryptone produced no PER, agreeing with previous work that the legs do not detect AAs for feeding initiation (Fig. 1B)^29,41,42^. In contrast, labellar stimulation with tryptone produced clear appetitive responses, with higher concentrations eliciting a more substantial response (Fig. 1C). As olfactory activation via food volatiles can potentially generate^43^ or enhance proboscis extension^42^, we repeated the labellar PER in flies with the antennae and maxillary palps removed, and in *Orco* mutants^44^. Tryptone PER was not significantly affected by the loss of olfaction via either method (Fig. 1 E-D). These results indicate that the AAs in a tryptone solution led to proboscis extension due to labellar contact chemosensation.

### Sweet, IR94e, and bitter GRNs act in parallel to detect amino acids

Next, we determined which of the five defined populations of GRNs on the labellum drive AA taste behaviors. GRNs are located in sensilla, each containing a different combination of cell types^45,46^. There are two appetitive sets of labellar chemosensory cells, “sweet” and "water”^47–54^. Each sweet GRN expresses *Gustatory Receptor 64f* (*Gr64f*) and responds to sugars^47,49–51^ and other appetitive chemical stimuli^52–54^. Water GRNs express *Pickpocket 28* (*ppk28*) and generally respond to solutions with low osmolarity^48,55,56^. The three remaining populations, “IR94e”, “bitter”, and “high salt” drive various levels of behavioral aversion. ^20,57–61^. Ionotropic Receptor 94e (IR94e) GRNs each express the *IR94e* gene, and we recently described their role in mediating mild feeding aversion to AAs^20^. Bitter GRNs express *Gr66a,* respond to a wide range of toxic stimuli^57–61^, and lead to aversive behaviors. *Ppk23* is expressed in high salt GRNs that specifically respond to concentrated salts to produce aversion that is dependent on internal salt homeostasis^45^. We used these genes to drive GRN-specific Gal4 expression to systematically determine which cell types respond to tryptone and contribute to external AA sensing.

Activation of sweet GRNs is sufficient to produce PER^20,62,63^, therefore, we hypothesized that sweet GRNs would drive the appetitive AA response. We used *Gr64f-Gal4* to drive the expression of the calcium indicator GCaMP7f and quantified the change in fluorescence at the GRN axon terminals in the subesophageal zone (SEZ) during tryptone stimulation. This *in vivo* imaging approach allows us to observe real-time calcium responses in taste cells that correlate with action potentials^20,64^. Sweet GRNs had significant responses to 5 and 10% tryptone compared to water, a negative control (Fig. 2A). To determine the behavioral role of sweet GRNs in tryptone PER, we used the optogenetic silencer GtACR1, a green-light-gated anion channelrhodopsin that requires all-*trans*-retinal (ATR) to function^65^. Tryptone PER was significantly decreased, but not eliminated, during sweet GRN silencing (Fig. 2B), which is consistent with their role in behavioral attraction. To determine if the appetitive water GRNs contribute to this response, we repeated these experiments with a *ppk28-Gal4* and found that low tryptone concentrations produced significant calcium responses (Fig. S1A). This result was expected, given that these GRNs respond to low osmolarity^46^. Optogenetic silencing of water GRNs during tryptone PER produced no change in behavior (Fig. S1B), suggesting that this cell type does not contribute to AA feeding, agreeing with previous work^18^.

**Figure 2:**
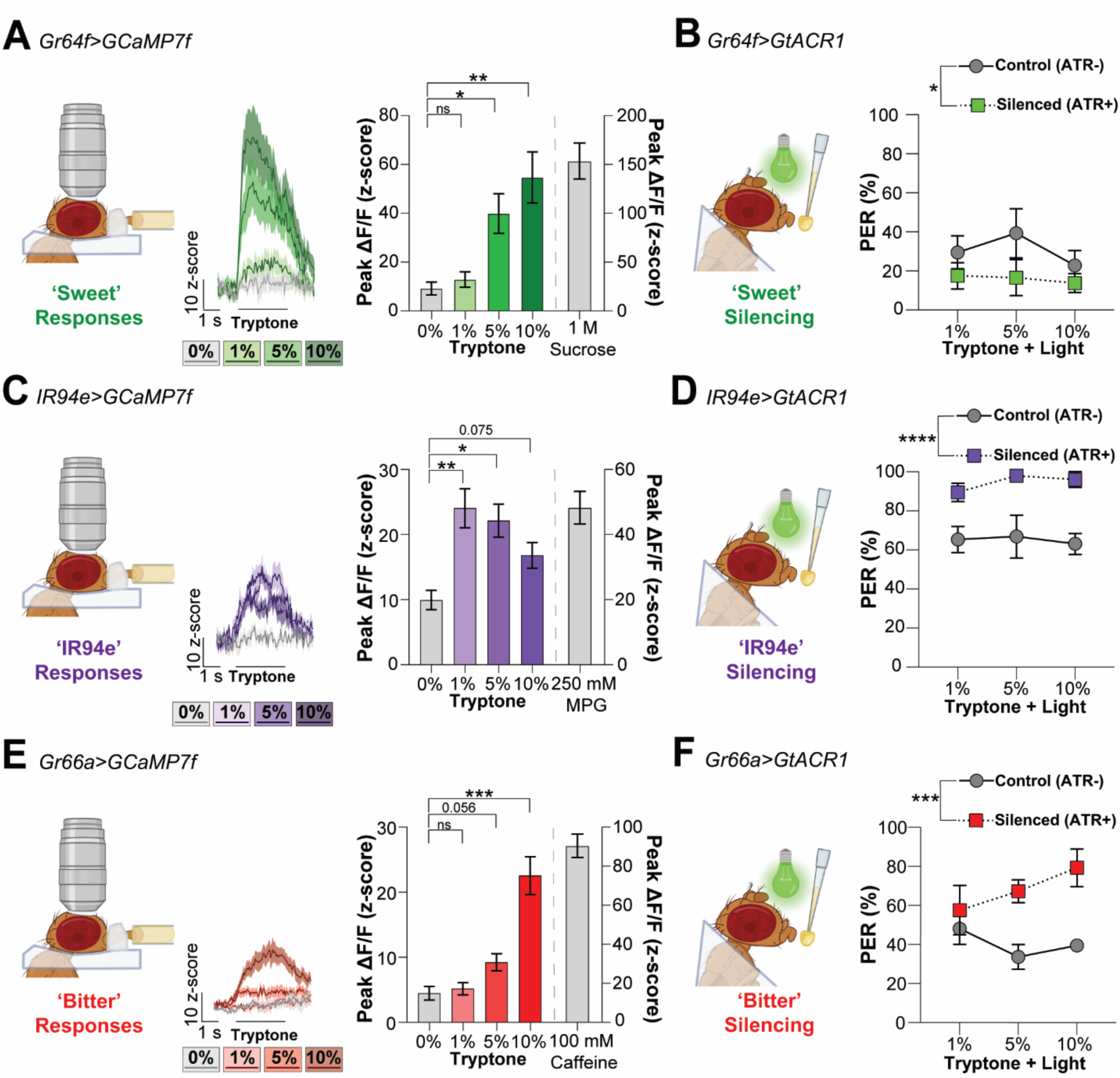
Tryptone detection is mediated through sweet, IR94e, and bitter GRNs. (**A, C, E**) *In vivo* calcium imaging in sweet (*Gr64f-Gal4*), IR94e (*IR94e-Gal4*), and bitter (*Gr66a-Gal4*) GRNs expressing *UAS-GCaMP7f*, n=13-15 flies. Fluorescence over time, bar indicates when stimulus is on the labellum (left). Peak fluorescence changes during the stimulation (right). Dashed line separates the axis change for the positive control in each population. (**B, D, F**) Tryptone PER in flies expressing *UAS-GtACR1* for optogenetic silencing in sweet (*Gr64f-Gal4*), IR94e (*IR94e-Gal4*), and bitter (*Gr66a-Gal4*) GRNs. n=5 groups of 5-10 flies. All data plotted as mean ± SEM. Trending p values indicated, ns=p>0.1, *p<0.05, **p<0.01, ***p<0.001, ****p<0.0001 by repeated measures one-way ANOVA with Dunnett’s posttest (A, C, E), or by two-way ANOVA (B, D, F).

To investigate the contribution of aversive GRNs, we started with IR94e because we recently found that these cells responded to solutions containing glutamate^6^. Using an *IR94e-Gal4* (*VT046252-Gal4*)^20,45^, we corroborated that 1 and 5% tryptone significantly increased calcium responses compared to the negative control (water), while 10% showed a strong trend approaching the cutoff for statistical significance (Fig. 2C). Behaviorally, silencing IR94e GRNs significantly increased PER to tryptone at all concentrations, suggesting that these GRNs contribute an aversive signal to limit AA PER (Fig. 2D). Using a *Gr66a-Gal4*, we found that bitter GRNs show concentration-specific calcium responses to tryptone: 1% elicited no visible responses, 5% produced a strong and visible trend, and 10% responses were statistically significant (Fig. 2E). Silencing bitter GRNs also significantly increased PER, particularly at high concentrations with a 40% increase to 10% tryptone (Fig. 2F). Repeating the imaging and behavioral experiments in high salt GRNs using a *ppk23-Gal4* indicated that they do not respond to any concentration of tryptone (Fig. S2C), nor do they impact tryptone PER (Fig. S2D), in agreement with previous findings^18^. In summary, sweet, IR94e, and bitter GRNs all contribute to the feeding initiation towards AAs. Our results suggest that sweet cells mediate attraction to AAs, whereas IR94e and bitter GRNs act to prevent AA feeding initiation, with low concentrations detected by IR94e cells and bitter GRNs responding to higher concentrations.

### AA feeding initiation is flexibly regulated through IR complexes and bitter GRs

To identify which receptors are necessary for tryptone detection in sweet, IR94e, and bitter GRNs, we first investigated the broadly expressed co-receptor, IR76b, which has been consistently implicated in AA detection^18,20,28,29,66^. *IR76b* mutants were used previously to establish this connection^18,28^, but we aimed to specifically knockdown *IR76b* in only one set of GRNs by expressing *IR76b* RNAi or an empty control using our GRN-specific drivers^67^.

Decreasing *IR76b* levels in sweet GRNs significantly reduced tryptone PER, with the largest changes occurring at 5 and 10% (Fig. 3A). We found a slight increase in PER when *IR76b* was knocked down in *IR94e* GRNs, but this did not reach statistical significance (Fig. 3C). This partially agrees with our previous finding that IR76b mutation abolished glutamate responses in IR94e GRNs^20^, although the knockdown is likely less efficient. *IR76b* knockdown in bitter GRNs led to a significant increase in PER (Fig. 3E), highlighting the role of this receptor in AA aversion.

**Figure 3:**
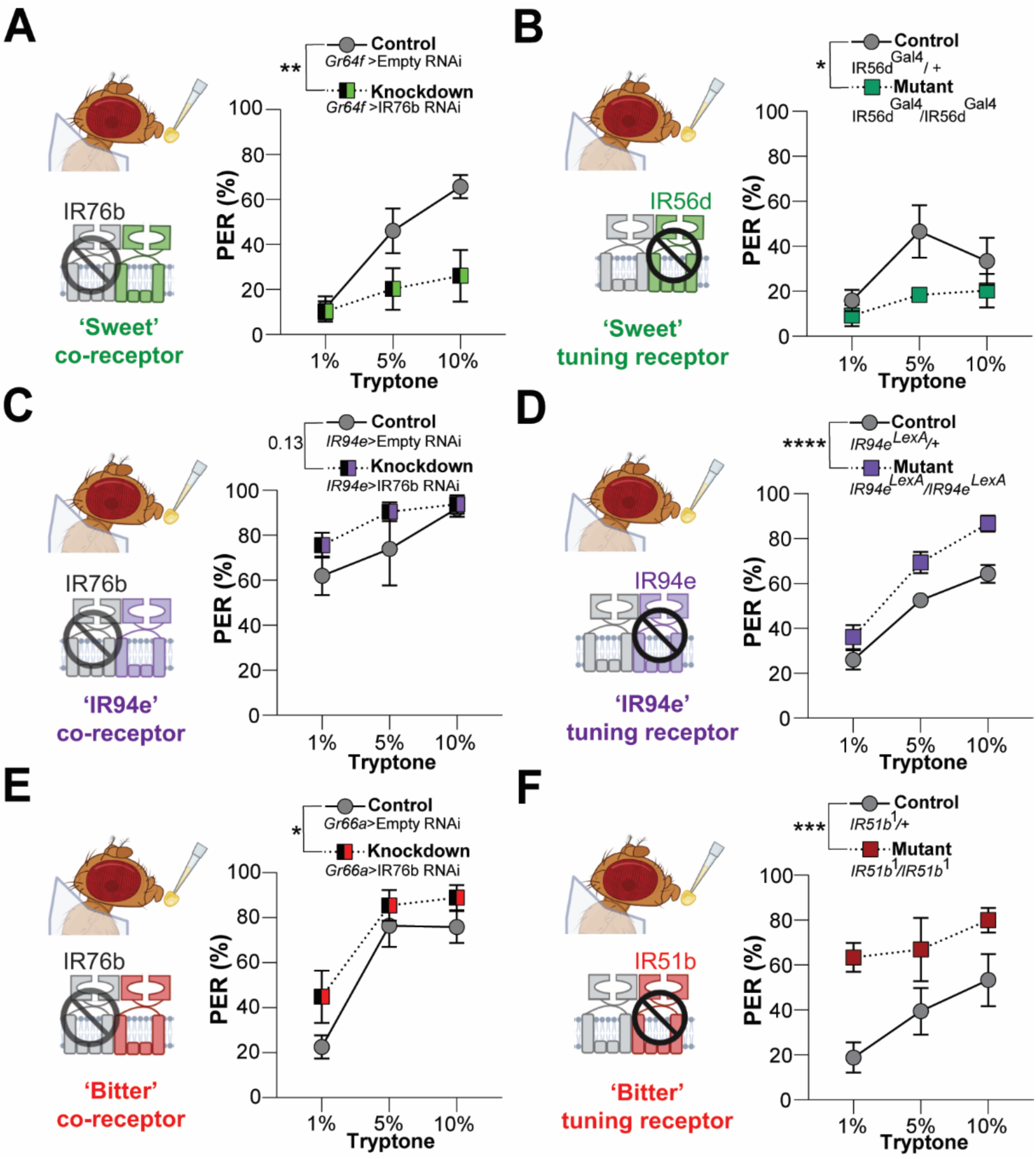
Ionotropic receptors contribute to labellar AA detection across GRNs. (**A, C, E**) PER to tryptone in flies expressing RNAi knockdown of the IR76b co-receptor in sweet (*Gr64f-Gal4*), IR94e (*IR94e-Gal4*), and bitter (*Gr66a-Gal4*) GRNs. (**B, D, F**) Tryptone PER in flies with mutations in tuning IRs within sweet (*IR56d*), IR94e (*IR94e*), and bitter (*IR51b*) GRNs. All data plotted as mean ± SEM. n= 5-6 groups of 5-10 flies for each genotype. Trending p values indicated, ns=p>0.1, *p<0.05, **p<0.01, ***p<0.001, ****p<0.0001 by two-way ANOVA.

Since *IR76b* knockdown in each type of GRN produced behavioral changes in the same direction as the cell silencing experiments, we screened for narrowly expressed tuning IRs that may form a complex with IR76b^66^. In sweet GRNs, we tested two candidate IRs known to be expressed in this population: IR56b, a low salt receptor^68^, and IR56d, a medium-chain fatty acid receptor^54,69,70^. *IR56b* mutants had no reduction in tryptone PER; instead, they had an unexpected increase (Fig S2A). In contrast, *IR56d* mutants showed a significant decrease in tryptone PER compared to heterozygous controls (Fig. 3B). While the PER to tryptone was not completely abolished with *IR56d* mutation, comparable to sweet GRN silencing, this result does suggest that IR56d mediates appetitive AA taste. Previous work identified IR94e as a tuning receptor for AAs, and in line with this role, PER to tryptone was significantly increased in *IR94e* mutants (Fig. 3D). Within bitter GRNs, IR51b was previously identified as a receptor for several individual AAs^18^, and we similarly found that *IR51b* mutation led to a significant increase in tryptone PER (Fig. 3F). Our results expand on IR76b, IR94e, and IR51b act as AA receptors and present a novel role for IR56d in detecting AAs.

Chemosensation through insect IRs mediates the detection of a broad spectrum of ligands^18,20,28,29,54,68–71^, and some tastants act through a combination of IRs and GRs^18,36^. In recent experiments, sugar-sensing Gr5a, Gr61a, and Gr64f were found to be involved in detecting low concentrations of certain AAs alongside IRs^18^. To determine if any receptors within the *Gr64* gene cluster contributed to tryptone detection, we tested flies with deletion of Gr*64a-f*^47^. Surprisingly, *ΔGr64a-f* flies had a significantly increased tryptone PER (Fig. 4A) despite a minimal PER to 1 M sucrose, confirming the absence of sugar receptors (Fig. S2D). This pattern of enhanced PER was consistent with the IR56b results (Fig. S2A) and persisted with individual sugar-sensing GR mutants: *Gr64f* mutation alone significantly increased tryptone PER (Fig. S2B) and *Gr5a* mutation mildly increased PER, although not to the level of statistical significance (Fig. 4B). Since these experiments produced phenotypes opposite to what we expected from the sweet cell silencing (Fig. 2B), we wanted to determine if this was a direct effect of the mutation (e.g. increasing sweet GRN responses to tryptone) or an indirect effect of the mutation on metabolism (e.g. mutants have altered hunger/satiety levels and show stronger PER due to an internal change). We used *in vivo* calcium imaging to confirm that this is a direct effect: the tryptone response in sweet GRNs of *ΔGr64a-f* mutants was significantly increased (Fig. 4C) despite the expected loss of sucrose responses (Fig. S2E). We hypothesized that this unexpected phenotype could be due to a change in the membrane space available for other receptors when the GRs are deleted from the genome. To determine if another IR-mediated taste modality in sweet GRNs was enhanced in these mutants, we tested PER to NaCl^68^. *ΔGr64a-f* mutants also had a significant increase in salt PER (Fig. S2C), suggesting that this is a nonspecific effect likely due to the presence of more IRs for detecting both salt and AAs when other receptors are absent.

**Figure 4:**
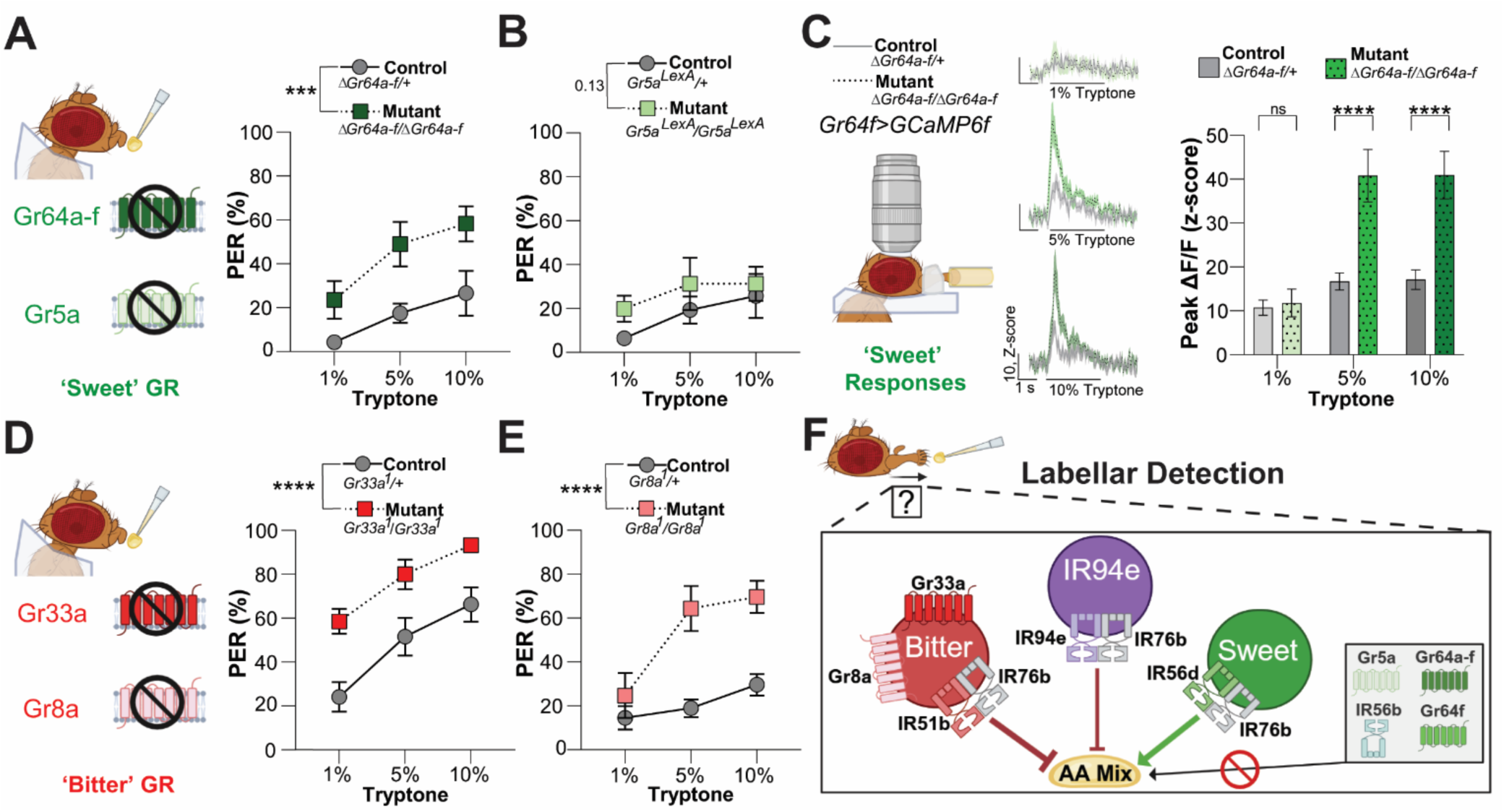
Bitter GRs contribute to AA detection. (**A**,**B**) Tryptone PER in flies with sugar-sensing GRs (*ΔGr64a-f*) and (*Gr5a*) mutations. n= 5-6 groups of 5-10 flies for each genotype. (**C**) Sweet GRN (*Gr64f-Gal4*) calcium imaging of GCaMP6f in sugar-sensing GR mutants (*ΔGr64a-f*). Fluorescence over time, bar indicates stimulus application (left). Peak fluorescence changes (right), n=11 flies per genotype. (**D**,**E**) Tryptone PER in flies with mutations in bitter-sensing GRs (*Gr33a*) and (*Gr8a*). n= 5-6 groups of 5-10 flies for each genotype. All data plotted as mean ± SEM. ns=p>0.05, *p<0.05, **p<0.01, ***p<0.001, ****p<0.0001 by two-way ANOVA (A,B,D,E) and two-way ANOVA with Šídák’s posttest (C). (**F**) Current model of the mechanisms behind labellar AA detection and feeding initiation.

A previous study found that bitter GRs were not involved in AA intake^18^, but we tested flies with a mutation in the broadly expressed bitter Gr33a receptor^72,73^ to confirm whether this is consistent with our tryptone mixture and feeding initiation. *Gr33a* mutants had a significant increase in PER (Fig. 4D), following the same pattern as bitter GRN silencing (Fig. 2F). To confirm this result, we tested a bitter-sensing GR that is only expressed in a subset of bitter GRNs, Gr8a^74,75^, which senses the non-proteinogenic AA L-canavanine^75^. Similarly, tryptone PER was significantly increased in *Gr8a* mutants (Fig. 4E). These results identify proteinogenic AAs as a new ligand for bitter GRs and suggest that a diverse set of IRs and GRs in bitter GRNs act redundantly to limit AA feeding.

## DISCUSSION

AA feeding must be tightly regulated to maintain homeostasis over an organism’s lifetime to improve the chances of survival^1–4^. We show that the detection and feeding initiation to a physiological mixture of AAs is regulated by sweet, IR94e, and bitter GRNs with unique combinations of IRs and GRs to allow for a flexible balance between attraction and avoidance. We confirm and expand upon the previously published roles of IR76b^18,20,28,29,66^, IR94e^20^, and IR51b^18^, redefining their role to include feeding initiation to a complex mix of AAs. Additionally, we identified that proteinogenic AAs also function as a ligand for IR56d, Gr33a, and Gr8a receptors (Fig. 4F).

### Amino acids activate an appetitive taste pathway

‘Umami’ taste is an AA cue in mammals that generally elicits feeding attraction^76^, and we find that the appetitive PER behavior in *Drosophila* is mediated through IRs in sweet taste cells. Using *in vivo* calcium imaging, we found that sweet GRNs responded strongly to AAs, but previous experiments using electrophysiological tip recordings to study sweet cell activation have produced varied results. Individual AAs or a small subset of AAs did not activate sweet GRNs in two studies^18,50,77^, whereas another study found that eight individual AAs significantly activated sweet cells via IR76b, IR25a, and sugar-sensing GRs^18^. While our IR76b results agree with the latter study, we found the opposite effect for GRs: sugar-sensing GR mutations increased neuronal and behavioral responses to AAs. The differences could be attributed to the solutions used, as previous work focused on individual AAs, and we used a mixture of AAs^18^.

Interestingly, a different study found that AA electrophysiological responses in L-type sensilla appeared only after a sugar-enriched but low-protein diet^78^, and AA deprivation can also enhance feeding initiation to promote the intake of nutrients that are lacking^29–31,41^. A similar behavioral phenomenon has been observed in humans, known as the protein leverage hypothesis, where the loss of dietary protein increases the intake of energy-dense foods to compensate for the missing nutrients^79^. Thus, AAs stimulate appetitive taste pathways across animals to encourage consumption, depending on nutrient needs.

Interestingly, the salience of AAs for feeding initiation is relatively weak compared to other nutrient sources like sugars^51^. However, the attraction to AAs is never entirely lost with our sweet cell silencing or sweet receptor manipulations. This implies that multi-sensory compensation may occur in the absence of attractive gustatory inputs, perhaps through olfaction, to allow for redundancy in detecting these essential nutrients. Similar compensation across sensory modalities has been described in *Aedes aegypti* mosquitoes^80^.

### Amino acid aversion is tightly regulated through redundant pathways

Though AAs are a necessary dietary component, excess intake can have detrimental effects across organisms^2–4^. We find that both IR94e and bitter cells act to regulate AA intake in *Drosophila*. Previous experiments have found that AAs consistently activate bitter cells through both electrophysiology and calcium imaging experiments, particularly at higher concentrations^18,59,77^. We corroborate the role of IR76b and IR51b^18^ as AA receptors in bitter cells and also find that bitter receptors Gr33a and Gr8a are involved in AA detection. In a previous study, bitter GR mutations did not affect arginine-specific electrophysiological responses^18^, suggesting that other AAs or a combination of AAs may act as ligands for these receptors.

IR94e GRNs have previously been studied with *in vivo* calcium imaging, where low Na+ and glutamate solutions act as ligands. ^20,45,81^. Here, we further confirm that IR94e GRNs respond to the glutamate-rich tryptone mixture through the IR94e receptor, particularly at lower concentrations. In a previous study, RNAi knockdown of *IR94e* had no impact on cellular activation or feeding preferences to arginine, serine, phenylalanine, or threonine^82^, suggesting that this cell type is more tuned to other AAs that are abundant in the tryptone AA mix, such as glutamate and aspartate^20^.

Finding that two aversive cell types mediate AA aversion through bitter GRs and multiple IRs emphasizes the importance of suppressing initial AA taste responses. This redundancy between receptor types is similar to heavy metal taste^36^, ultimately preventing the overconsumption of specific molecules. In comparison, it has been observed that multiple human bitter taste receptors can bind to phenylalanine and tryptophan to elicit cellular responses^13,83^. This observation highlights the unique dynamics occurring across cells and receptors when encountering a mix of AAs. In *Drosophila*, dietary AA balance may not be the only state to impact feeding behaviors, as IR94e cells also directly contribute to egg laying^20^. This leads to an interesting hypothesis that the aversive cells guiding behaviors for AAs at certain concentrations may be allowing flies to balance their dietary intake needs with the nutrient needs of their offspring.

### Amino acid taste mechanisms display multiple examples of biological convergence

AA detection across all three cell types involves IRs, establishing this type of receptor as one of *Drosophila’s* primary AA detectors. Though gustatory receptors between mammals and *Drosophila* differ^84^, there is some conservation between mammalian ionotropic glutamate receptors (iGluRs) and *Drosophila* IRs^85^. Even though the ligand binding domain of many IRs lacks key glutamate binding motifs^85^, it has been proposed that this evolutionary relationship may allow IRs to potentially function as AA receptors^86^, which we and others have now confirmed^18,20,28,29^. Furthermore, IR51b and IR56d AA sequences across *Drosophilid* species align into sister groups^86^, supporting our findings that both receptors detect similar AA ligands.

New dietary niches modify taste receptors across animals^87^, and in primates, switching from insectivorous to foliage-based protein diets containing higher concentrations of glutamate has led to repetitive receptor evolution towards glutamate sensitivity^88^. Classically, glutamate taste has been difficult to investigate as it comes in salt forms, which have the confound of activating salt taste pathways, or as glutamic acid, which has low solubility in water^89^. Tryptone is advantageous as it contains glutamate as the most abundant AA along with a mix of AAs common in foodstuffs, which we find interacts uniquely with taste cells and receptors. From mammals to insects, the ability of gustatory systems to detect abundant AAs in foodstuffs, such as glutamate, appears to be evolutionarily convergent.

In conclusion, this work is foundational to understanding how AA feeding is initiated through parallel gustatory inputs and advances our understanding of AA taste coding across animals. Future work can determine whether internal states act directly on these specific gustatory pathways to modulate AA taste sensitivity or protein intake.

### Limitations

As highlighted above, AA feeding is multi-faceted. Our utilization of hungry mated females does not encompass all the possible behaviors toward AAs. Our goal was to create a basic map of the AA taste mechanisms in *Drosophila* for future study. Additionally, our assays describe feeding initiation to tryptone-based AA mixtures, which is biologically relevant, but a given individual AA may not work through all of the cells and receptors identified here. Finally, our study utilized PER to specifically isolate initial AA detection and avoid any internal AA-sensing mechanisms that could also impact AA feeding. Therefore, we described AA detection via the external taste organs only, whereas other studies have identified mechanisms of AA detection using preference or intake assays, where AAs can activate pharyngeal taste cells^90^ and prolonged feeding events can alter AA homeostasis and taste sensitivity^29^.

## ACKNOWLEDGEMENTS

We thank the Bloomington *Drosophila* Stock Center (BDSC) and Dr. Michael Gordon for fly stocks. This work was supported by NSF award #2332375 and funds from the Department of Biology and the College of Arts and Sciences at the University of Vermont. Graphics were generated in BioRender.com.

## AUTHOR CONTRIBUTIONS

M.S. conceptualized the project, secured funding, and provided supervision. J.G. and G.D. wrote the original manuscript and performed all formal data analysis. J.G. produced all the figures. All authors contributed to data collection and reviewed the manuscript.

## DATA AVAILABILITY

A Mendeley dataset containing the original data generated in this study will be made publicly available upon formal acceptance and linked to this manuscript.

---

**Supplemental Figure 1:**
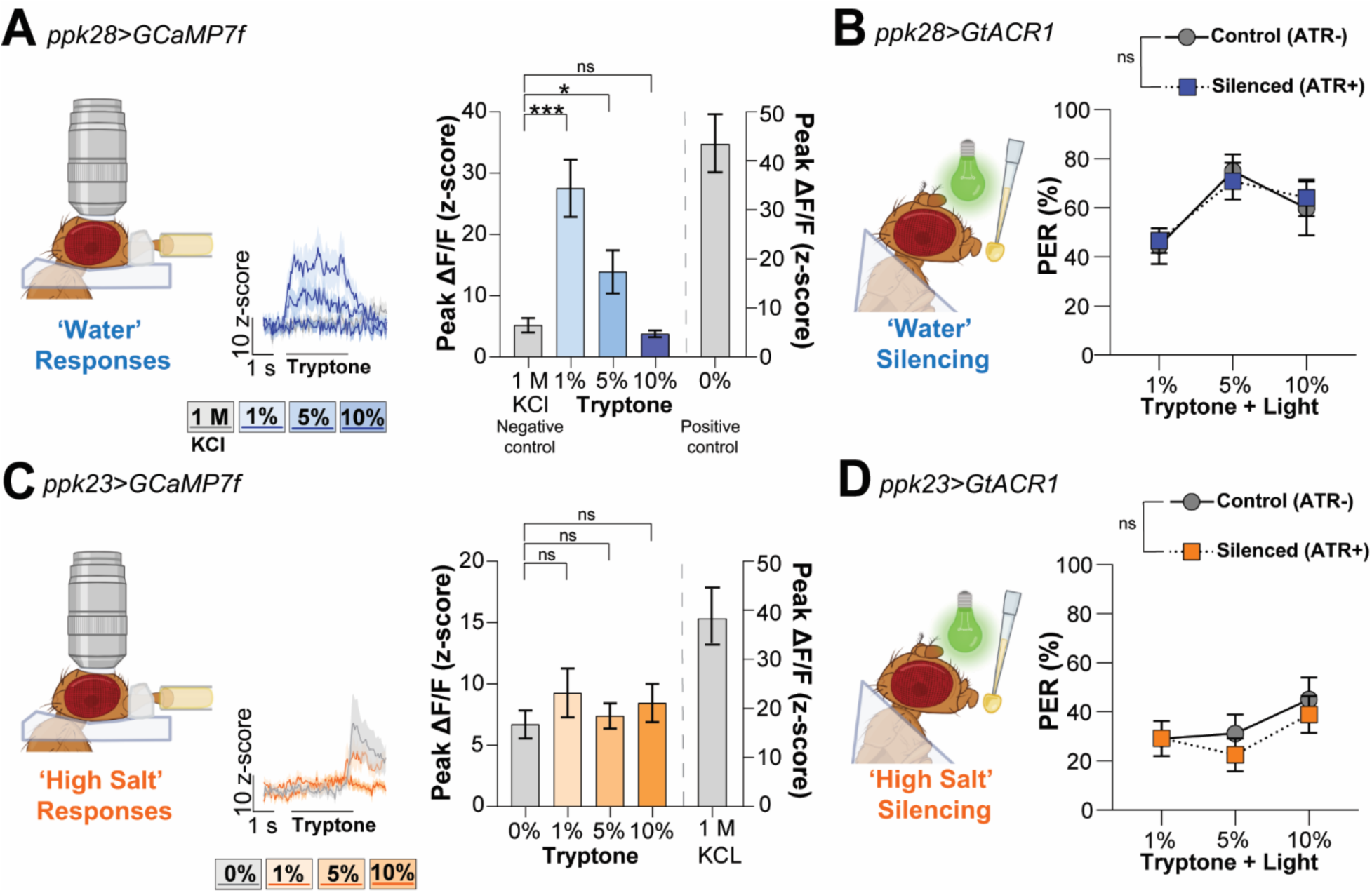
Water and high salt GRNs do not contribute to tryptone taste responses. Related to Figure 2. (**A,C**) Calcium imaging during tryptone stimulation in water (*ppk28-Gal4*) and high salt (*ppk23-Gal4*) GRNs driving *UAS-GCaMP7f,* n=12-13 flies. Change in fluorescence over time, bar indicates when stimulus is on the labellum (left). Note: low osmolarity solutions generate a small OFF response with tastant removal (C). Peak fluorescence changes during the stimulation (right). Dashed line separates the axis change for the positive control in each population. (**B,D**) PER to tryptone during optogenetic silencing via GtACR1 in water (*ppk28-Gal4*) and high salt (*ppk23-Gal4*) GRNs. n= 5 groups of 5-10 flies each. All data plotted as mean ± SEM. ns=p>0.05, *p<0.05, ***p<0.001 by repeated measures one-way ANOVA with Dunnett’s multiple comparisons (A,C), or by two-way ANOVA (B,D).

**Supplemental Figure 2:**
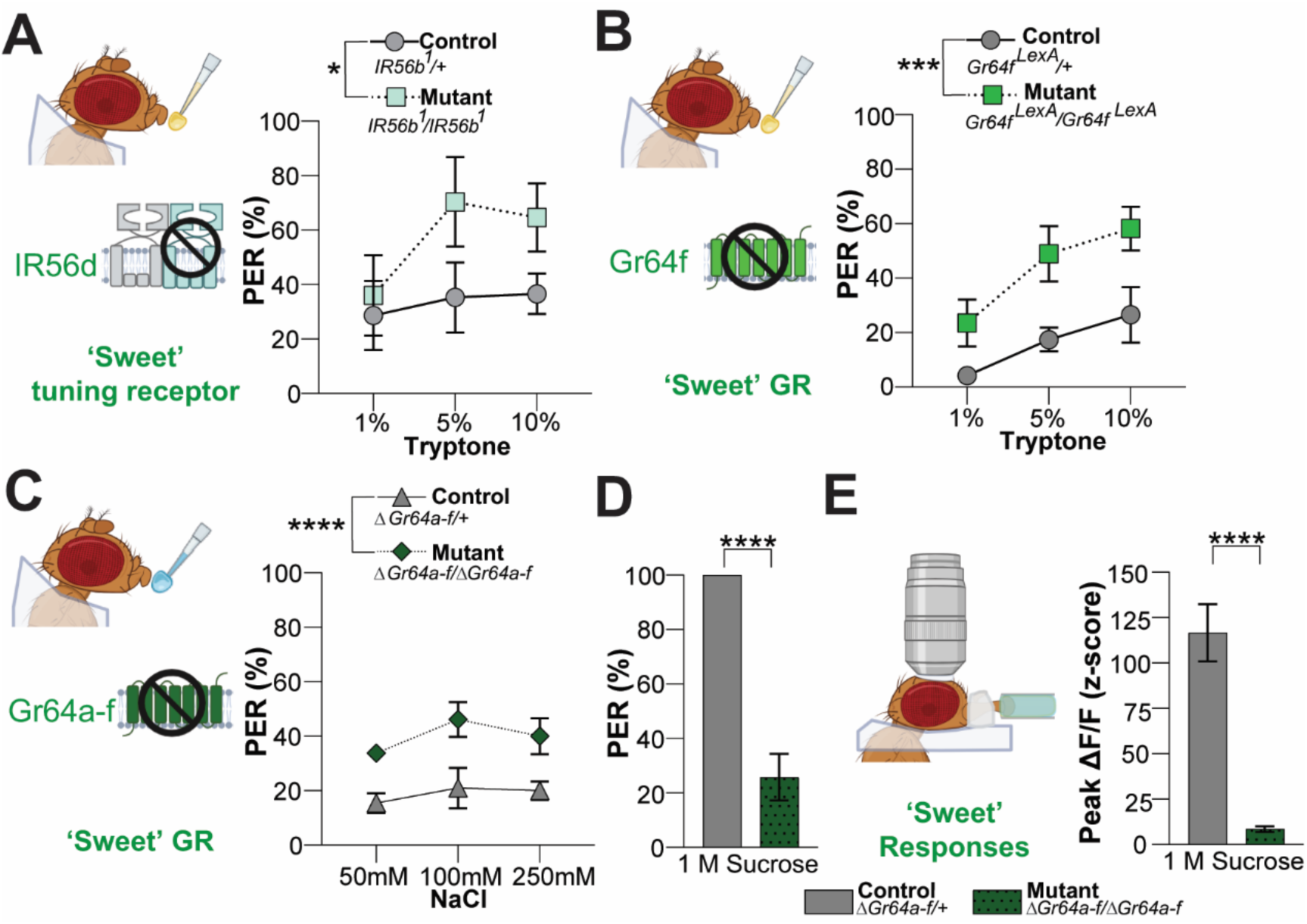
Receptor mutations in sweet GRNs lead to enhanced responses despite a loss in sugar sensing. Related to Figure 4. **(A)** Tryptone PER in another tuning IR in sweet GRNs (*IR56b*). (**B)** Tryptone PER in a single receptor within the *Gr64* cluster (*Gr64f*). (**C**) PER to salt in the *Gr64* cluster mutants (*ΔGr64a-f*). (**D**) PER to sucrose in the *Gr64* cluster mutants (*ΔGr64a-f*). n=5-6 groups of 5-10 flies for each genotype. (**E**) Peak fluorescence change to 1 M sucrose in *Gr64f>GCaMP6f* flies during calcium imaging of sweet GRNs with the *Gr64* cluster mutation (*ΔGr64a-f*), n=11 flies per genotype. All data plotted as mean ± SEM. *p<0.05, ***p<0.001, ****p<0.0001 by two-way ANOVA (A-C), and Student’s t-test (E-D).

## METHODS

### Key resources table

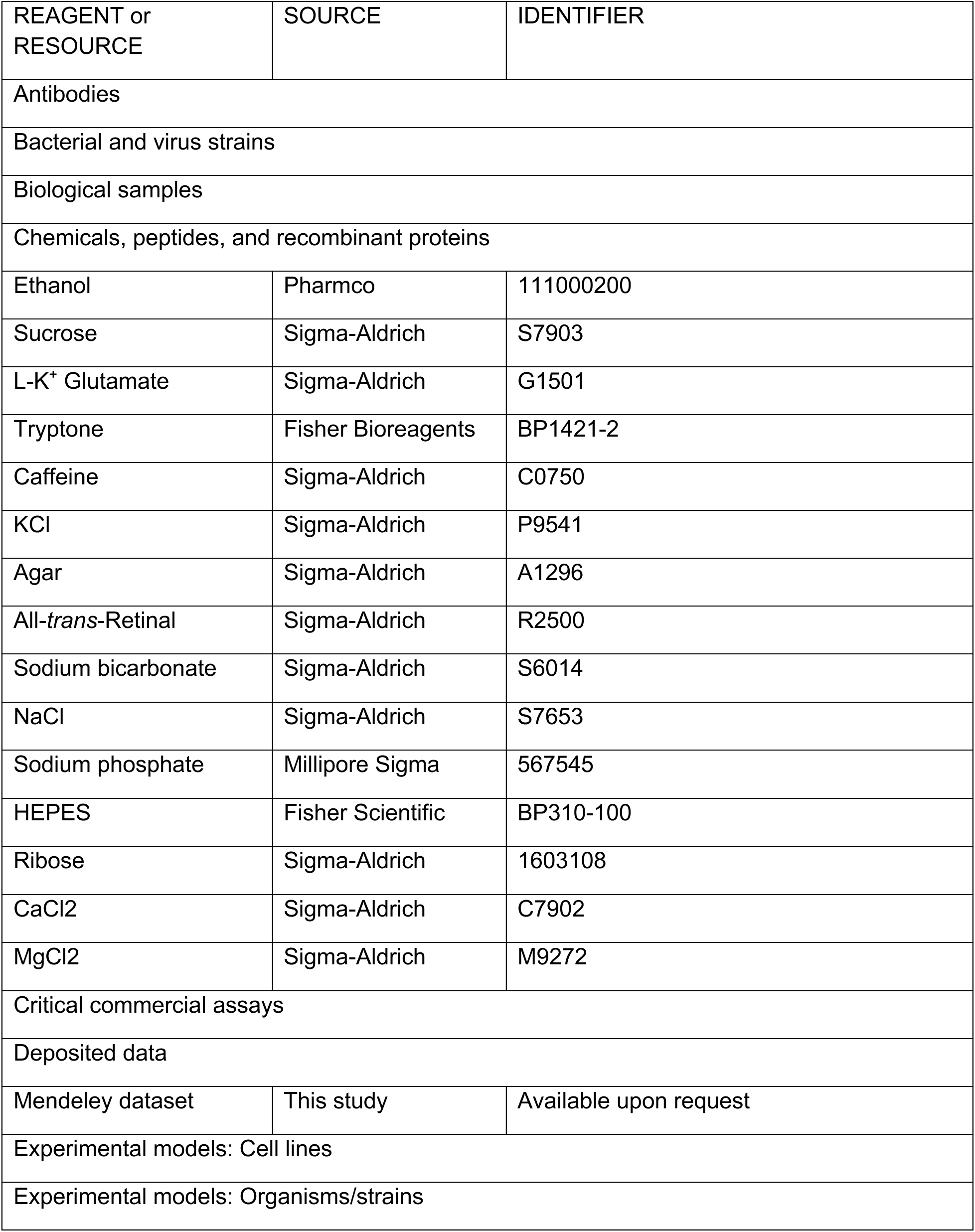
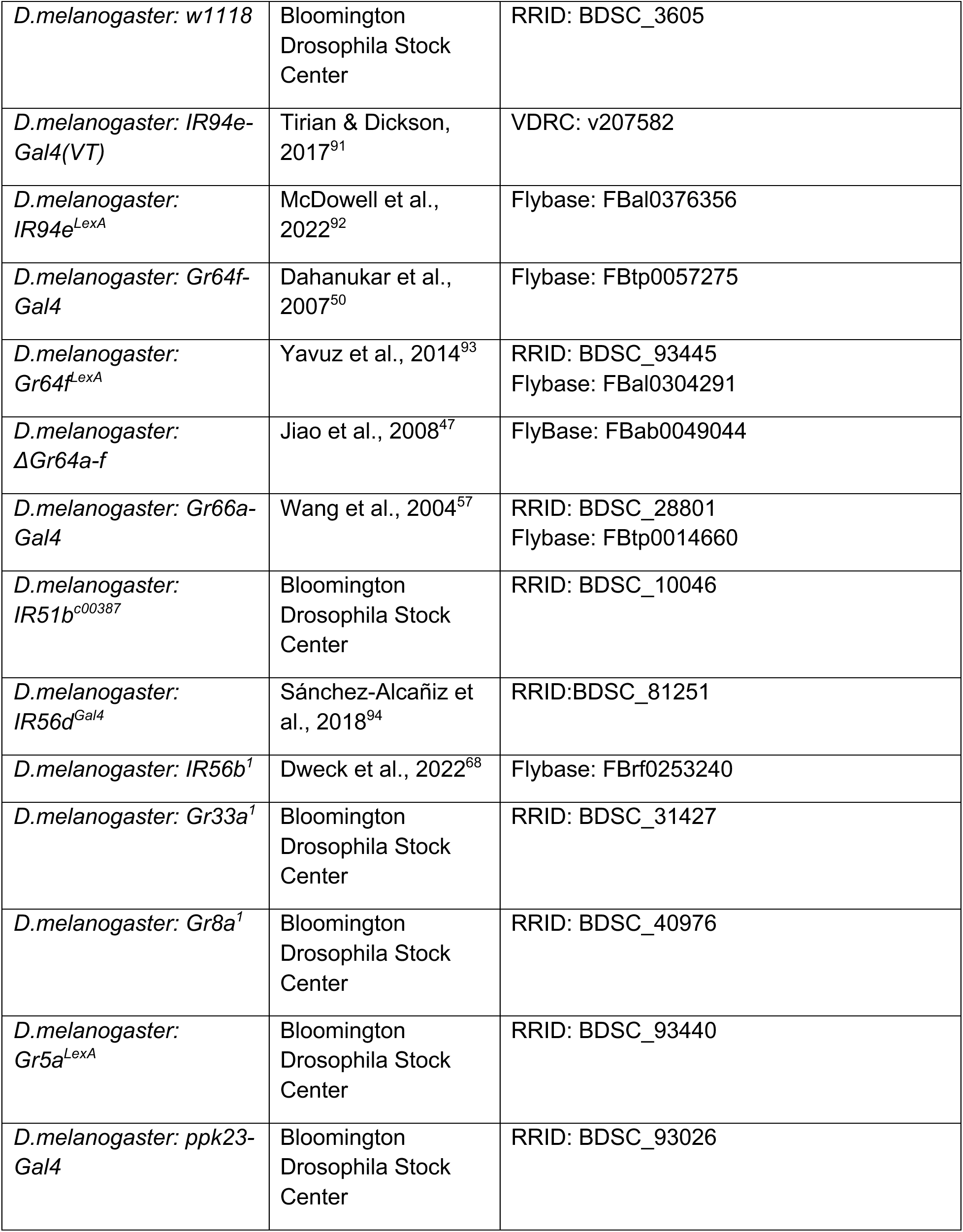
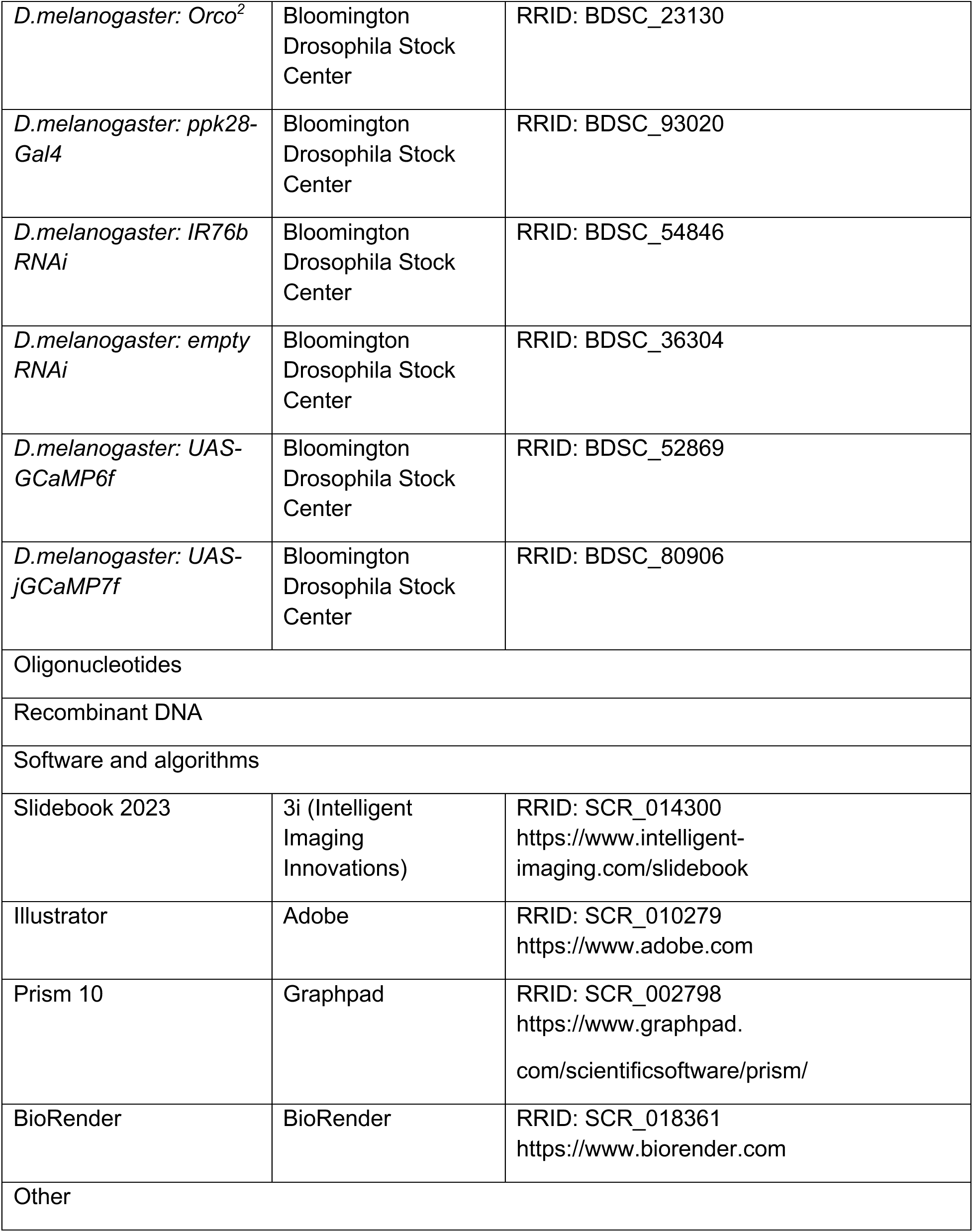

### Lead contact

Additional information and requests for resources and reagents should be directed to and will be fulfilled by the lead contact, Molly Stanley.

### Flies

Experimental flies were kept on regular cornmeal food at 25°C in 60% relative humidity. Mated female flies between 2-10 days old were used in all experiments. The fly stocks used for experiments are listed in the Key Resource Table and genotypes are indicated in each figure.

### Chemicals

A full list of chemicals with source information can be found in the Key Resources Table. Sucrose and K^+^ glutamate were dissolved in water at specified concentrations. Tryptone was freshly made up in water at the indicated w/v% solutions. All-*trans*-retinal (ATR) was made up in 100% EtOH, kept at -20°C, and diluted to a final concentration of 1 mM with EtOH of the same dilution given as a vehicle.

### Amino acid analysis

All AA data were curated from available databases and literature. AA content from apples, bananas, grapes, and tomatoes were collected directly from the United States Department of Agriculture FoodData Central database. All samples were selected to be in their pure form and concentrations were determined out of 100g of each food^27^. To determine the AAs present in tryptone, we curated data from the enzymatic hydrolytic digest of casein^38^, which is the basis of tryptone. We chose 27.6% degrees of hydrolysis as our reference point as it was likely the peak of the protein digestion and similar to the AA mixture found in commercially available tryptone^38^. To find the approximate AA concentrations of yeast extract we found the mean of each AA within the range of concentrations published^33^. For each source, the AA are plotted as the proportion of total AAs present.

### Behavioral assays

<u>Labellar PER:</u> 24 hours prior to the assay, flies were transferred to starvation vials containing 1% agar. Flies were mounted for the assay as previously described^20,39^. Briefly, a mouth pipette was used to transfer flies into 200 µL pipette tips cut so only the heads were exposed, permitted to recover in a humidity chamber for ∼1 hour, and water satiated. Water was presented as the first stimulus as a negative control (to ensure flies did not exhibit PER to water). Tryptone at increasing concentrations was presented and the final stimulus was 1 M sucrose, a positive control (to ensure that flies were able to exhibit PER). Flies that responded to H2O stimulation prior to stimulation with the experimental solutions were excluded. Flies that did not respond to 1 M sucrose after stimulation with the experimental solutions were excluded, except in the case of sugar GR mutants.

<u>Tarsal PER:</u> 24 hours prior to the assay, flies were transferred to starvation vials containing 1% agar. Flies were anesthetized and mounted on slides using quick-dry nail polish, ensuring immobilization while keeping the tarsi exposed. Mounted flies were labeled and placed in a humidity container for recovery for ∼1 hour and water satiated. Water was presented as the first stimulus as a negative control (to ensure flies did not exhibit PER to water). Tryptone at increasing concentrations was presented and the final stimulus was 1 M sucrose, a positive control (to ensure that flies were able to exhibit PER). Flies that responded to H2O stimulation prior to stimulation with the experimental solutions were excluded. Flies that did not respond to 1 M sucrose after stimulation with the experimental solutions were excluded.

<u>Optogenetic PER:</u> three days prior to the assay, flies were collected and placed on all-*trans*-retinal (ATR) or ethanol mixed into normal food for two days as previously described^20,65,95^. One day prior to the assay, flies were transferred to starvation vials containing ATR or ethanol mixed into 1% agar. Vials were covered with aluminum foil to minimize exposure to light and kept at 25°C for 24 hours. Flies were mounted for the aforementioned labellar PER assay. Flies were mounted in a dark room with minimal light under a dissection microscope. Water was presented as the first stimulus as a negative control (to ensure flies did not exhibit PER to water). The experimental tryptone solutions were presented in combination with a green LED powered by a 9V battery (∼425 µWatts), held directly over the fly labellum. The final stimulus was 1 M sucrose, a positive control to ensure that flies were able to exhibit PER. The efficacy of optogenetic inhibition was validated by confirming the complete suppression of PER to 1 M sucrose in *Gr64f>*Gt*ACR1*, ATR+ flies in the presence of the green LED (data not shown). Flies that responded to H2O stimulation prior to stimulation with the experimental solutions were excluded. Flies that did not respond to 1 M sucrose after stimulation with the experimental solutions were excluded.

### Calcium imaging

*In vivo* imaging of labellar GRN axon terminals was performed as previously described^20,96^. First, mated female flies were anesthetized with CO2 and mounted into a specialized chamber in which their heads are secured with nail polish. Once secured, the fly’s labellum was manually extended and waxed into this position to ensure that the proboscis was unobstructed. Flies were then permitted to recover in a humidity chamber for 1 hour. After recovery, cuticle above the subesophageal zone (the imaging area) was dissected to expose the brain. While the brain was exposed, adult hemolymph-like (AHL) solution (108 mM NaCl, 5 mM KCl, 4 mM NaHCO3, 1 mM NaH2PO4, 5 mM HEPES, 15 mM ribose, 2mM Ca2+, 8.2mM Mg2+, pH 7.5) was continuously applied to the brain. The imaging area was cleared of obstructions by cutting and removing the respiratory tissues and part of the esophagus. Flies were imaged with a 3i spinning disc confocal station (Zeiss upright microscope, 2Kx2K 40 fps sCMOS camera, CSU-W1 T1 50 mm spinning disc). AHL was used as the immersive solution during image acquisition with a 40x water immersion objective and 2x zoom. Once positioned for imaging, baseline fluorescence was recorded before the fly’s labellum was stimulated with the tastant for 5 seconds, and the recording continued for five seconds after stimulus removal. The tastant was directly applied to the labellum via a micromanipulator and a capillary tube which fit directly over the fly’s labellum. Each fly was exposed to 5 total solutions, a negative control, 1,5,10% tryptone, and a positive control. The positive and negative controls differed depending on the neuronal population being imaged and are indicated in the figures. The camera was set to 100 ms exposure and 9.5 Hz capture rate. In Slidebook, background fluorescence was subtracted for each image and a region of interest was drawn around the GRN projections. The mean intensity was exported for further analysis. Ten points of baseline were averaged and delta F/F (%) for each point calculated as ((F - Baseline F) / Baseline F)*100). Three consecutive points with the highest peak change while the stimulus was over the labellum, during the stimulus onset, prior to the removal of the stimulus, were averaged to reflect the peak response. Flies were excluded if they showed a large response to the negative control (>30%) or no response to the positive control (<15%).

### Quantification and statistical analysis

All statistical tests were performed in GraphPad Prism 10 software. Specific tests are indicated in the figure legends along with the number of replicates. Experimental or genotype controls were always run in parallel. Behavioral assays were repeated across multiple days and genetic crosses. Data are plotted as mean +/- SEM in bar graphs and line graphs. Asterisks indicate *p<.05, **p<.01, ***p<.001, ****p<.0001.

